# Viral infection impacts volatile organic compound production in the coccolithophore *Gephyrocapsa huxleyi*

**DOI:** 10.1101/2025.02.09.637333

**Authors:** Holger H. Buchholz, Kristina D.A. Mojica, Jacqueline Comstock, Lindsay Collart, Stephen J. Giovannoni, Kimberly H. Halsey

**Affiliations:** Department of Microbiology, Oregon State University, Corvallis, OR, USA; School of Ocean Science & engineering, University of Southern Mississippi, MS, USA; Marine Science Institute, University of Southern California, Santa Barbara, CA, USA

## Abstract

Volatile Organic Compounds (VOCs) are a diverse collection of molecules critical to cell metabolism, food web interactions, and atmospheric chemistry. The eukaryotic coccolithophore *Gephyrocapsa huxleyi*, an abundant coastal eukaryotic phytoplankter, forms massive blooms in coastal upwelling regions, which are often terminated by viruses (EhVs). *G. huxleyi* produces organosulfur VOCs such as dimethyl sulfide (DMS) and halogenated metabolites that play key roles in atmospheric chemistry. Here we resolved the role of lytic viral infection by EhV207 on VOC production of the model strain *G. huxleyi* CCMP374. Our analysis identified 79 VOCs significantly impacted by viral infection, particularly during cell lysis, with sulfur containing VOCs like DMS dominating the profiles. Viral lysis results in a nearly six-fold increase in VOC production and generated a previously unrecognized range of VOCs, including 15 sulfur, 22 nitrogen, 2 phosphorus, 19 oxygen and 17 halogen-containing compounds. These findings reveal that viral infection of *G. huxleyi* releases VOCs which are much more diverse than previously recognized. We further show that EhV207 primarily accelerates existing metabolic processes in *G. huxleyi* and facilitates the release of pre-existing intracellular VOCs rather than inducing novel biochemical pathways. This wide range of VOCs may be produced on a massive scale during coccolithophore bloom-and-bust cycles, with important impacts on coastal biogeochemistry and surface ocean/atmosphere interactions.

## Introduction

Volatile Organic Compounds (VOCs) are the evaporative fraction of the dissolved organic carbon (DOC) pool and constitute a diverse group of chemicals that include a large range of climate-active gases. In the environment, the cosmopolitan coccolithophore *Gephyrocapsa huxleyi* (formerly *Emiliania huxleyi*) (Bendif *et al*., 2019) is known to produce many oxygenated VOCs such as methanol, acetaldehyde, isoprene and the sulphur containing VOCs, dimethyl sulfide (DMS) and DMSP, as well as halogenated VOCs (Sinha *et al*., 2007; Colomb *et al*., 2008; Mincer and Aicher, 2016; Yu *et al*., 2022). Oceanic emission is a major source of atmospheric VOCs and algal blooms, hotspots of primary production in the ocean, are thought to fuel the VOC pool. The total VOC pools are reduced via oxidization by heterotrophic cells (Moore *et al*., 2020, 2022), making VOCs a significant player in the carbon cycle of the surface ocean. A major driver of phytoplankton bloom demise is lytic viral infection (Bratbak *et al*., 1996; Fuhrman, 1999), wherein cell lysis leads to the liberation of carbon from biomass to the DOC pool, a process referred to as the “viral shunt” (Suttle *et al*., 1990; Wilhelm and Suttle, 1999). This ecological process reduces energy transfer to higher trophic levels and plays key roles in nutrient cycles of marine microbial communities. However, it is largely unknown how algal bloom succession shapes VOC composition and how viruses affect the VOC pool.

DMS and other organic sulfur volatiles are recognized as major VOCs and components of the global biogeochemical sulfur cycle (Kettle and Andreae, 2000; Moran and Durham, 2019). Especially in cold water environments, *G. huxleyi* and other phytoplankton cause large amounts of DMS emission to the atmosphere and can account for up to 70% of oceanic sulfur emissions (Simon *et al*., 2009; Carpenter and Bennett, 2011; Dani and Loreto, 2017). Atmospheric DMS may contribute to cloud condensation nuclei over the oceans, though the extend of which is debated (Quinn and Bates, 2011). This has been proposed to influence climate feedbacks associated with temperature-induced DMS production by phytoplankton (Charlson *et al*., 1987). The ocean is also a major source of reactive atmospheric halogens like chlorine, bromine and fluorine (Solomon et al., 1994). Bromoform (CHBr_3_), for example, has a significant role in stratospheric ozone destruction (Anbar et al., 1996; Barrie et al., 1988). In one of the first studies investigating halogenated VOC emissions from phytoplankton, Colomb et al. (Colomb *et al*., 2008) suggest that *G. huxleyi* is a weak emitter of halogens compared to other taxa that produce halogenated terpenes, oxylipins, polyketides, bromomethane and bromoform, and iodocarbons (Moore and Tokarczyk, 1993; Manley and de la Cuesta, 1997; Scarratt and Moore, 1998; Laturnus, 2001; Colomb *et al*., 2008; Cabrita *et al*., 2010; Paul and Pohnert, 2011). Yet, when coccolithophore blooms are terminated by viral infection, they leave a distinct footprint of halogenated DOM compounds (Kuhlisch *et al*., 2021), implying that coccolithophore bloom succession is a driver of marine halogenated VOC cycles.

Despite the importance of VOCs released by algae and viral lysis, and their role in atmospheric VOC chemistry and climate feedbacks, they are frequently overlooked in marine carbon studies. VOCs are part of the labile DOC pool, to which estimates suggest viral lysis and senescent cells contribute 40% (Moran *et al*., 2022). Blooms of *G. huxleyi* and their termination by giant double stranded DNA viruses (EhVs) have been well studied and are an important model host-virus system. Viral hijacking of *G. huxleyi* hosts leads to a profound rewiring of metabolic pathways to support viral replication. EhV infection can increase the release of DMS, acrylic acid, halogenated metabolites and reactive oxygen species (ROS) (Evans, Malin, Mills, *et al*., 2006; Kuhlisch *et al*., 2021). They furthermore promote transparent exopolymer particle (TEP), particulate inorganic carbon (PIC) and DOC release; and thereby EhVs impact dynamics of heterotrophic microbial communities that are interacting with these released compounds (Evans, Malin, Mills, *et al*., 2006; Evans *et al*., 2007; Joassin *et al*., 2011; Sheyn *et al*., 2016; Vincent *et al*., 2023). However, there is a disconnect between the diversity of VOCs observed in the environment and our knowledge of their production by ecologically important algae such as *G. huxleyi*.

Understanding the microbial ecology of VOCs has been challenged by difficulties in accurately measuring VOCs at low concentration. This is primarily because it was difficult to measure many VOCs before appropriate instruments such as proton-transfer reaction mass spectrometry (PTR-MS) became available (Halsey *et al*., 2017). We coupled PTR-MS with environmentally controlled dynamic stripping chambers, which use VOC-free air passing through samples to direct VOCs dissolved in the sample into the PTR-MS for proton transfer (using H_3_O+ and O_2_+ as ion source) ^(^Davie-Marti^n^ *et al*., 2020; Moore *et al*., 2020). In this study, we investigated the impact of EhVs on the total VOC pool released from coccolithophores. Specifically, we infected *G. huxleyi* CCMP374 cultures with the model virus EhV207 and assessed total extracellular VOCs using PTR-MS and stripping chambers. We found viral infection to have a major impact on VOC production, altering the range and quantities of VOCs produced, and producing many VOCs with m/z values not previously recorded in coccolithophore metabolite profiles.

## Results and Discussion

### Growth characteristics of virally infected G. huxleyi

24 hours after infecting axenic cultures of *G. huxleyi* CCMP374 with lytic EhV207, cell concentrations decreased in infected cultures compared to uninfected controls (Fig. 1a-b). Declining cell concentrations were concomitant with an increase in viral concentration, indicating the onset of viral lysis. The fraction of cells with compromised membranes were elevated in infected cultures after 24h (10 %) compared to controls (<1 %) (Figure 1e). Flow cytometry on SYTOX stained infected cells during the incubation period confirmed that the cell membranes were not compromised from virion attachment and genome injection, in accord with recent reports that EhV201 (closely related to EhV207(Allen *et al*., 2007; Nissimov *et al*., 2012)) has an outer and inner membrane around the capsid that fuses with the host membrane to inject viral genetic material, rather than puncturing through the cell membrane (Mackinder *et al*., 2009; Homola *et al*., 2024). This could limit host cell membrane permeability that is sometimes associated with viral infection (Puck and Lee, 1955),Young et al. 2000, Sobhy 2017) and which could otherwise increase the rate of VOC diffusion through the membrane. After 48 hours, cells in the infected cultures had visibly cleared. Autofluorescence-based flow cytometry detected cellular signals that were likely dead or dying cells, coinciding with the rise of virus abundance (∼3×10^8^ viruses mL^-1^). Less than 1 % of cells had compromised membranes in uninfected controls at any time point, suggesting they consisted of healthy and active cells throughout the experiment even after reaching a maximum cell density of ca. 2×10^6^ cells mL^-1^ around 48-72 hours after the start of the experiment. This conclusion is supported by the steadily increasing fluorescence (F_0_) values and consistent F_v_/F_m_ (ca. 0.6) values observed. Conversely, F and F_v_/F_m_ of infected cultures decreased 24 hours post infection. In the following discussion of results, for a total of seven time points over 96 hours, the first 24 hours are considered the latent period; the period between 24 and 48 hours is the late infection stage, including the bulk of cell lysis; and timepoints between 48-96 h are post lysis. For each time point, the integrated sum of VOC concentrations (ppbv) for each peak (m/z) between masses m/z 30 to 350 (using separate sacrificial cultures for measurements with hydronium and ozone as ion sources) were combined for the analysis (Supplementary Figure 1).

**Figure 1.**
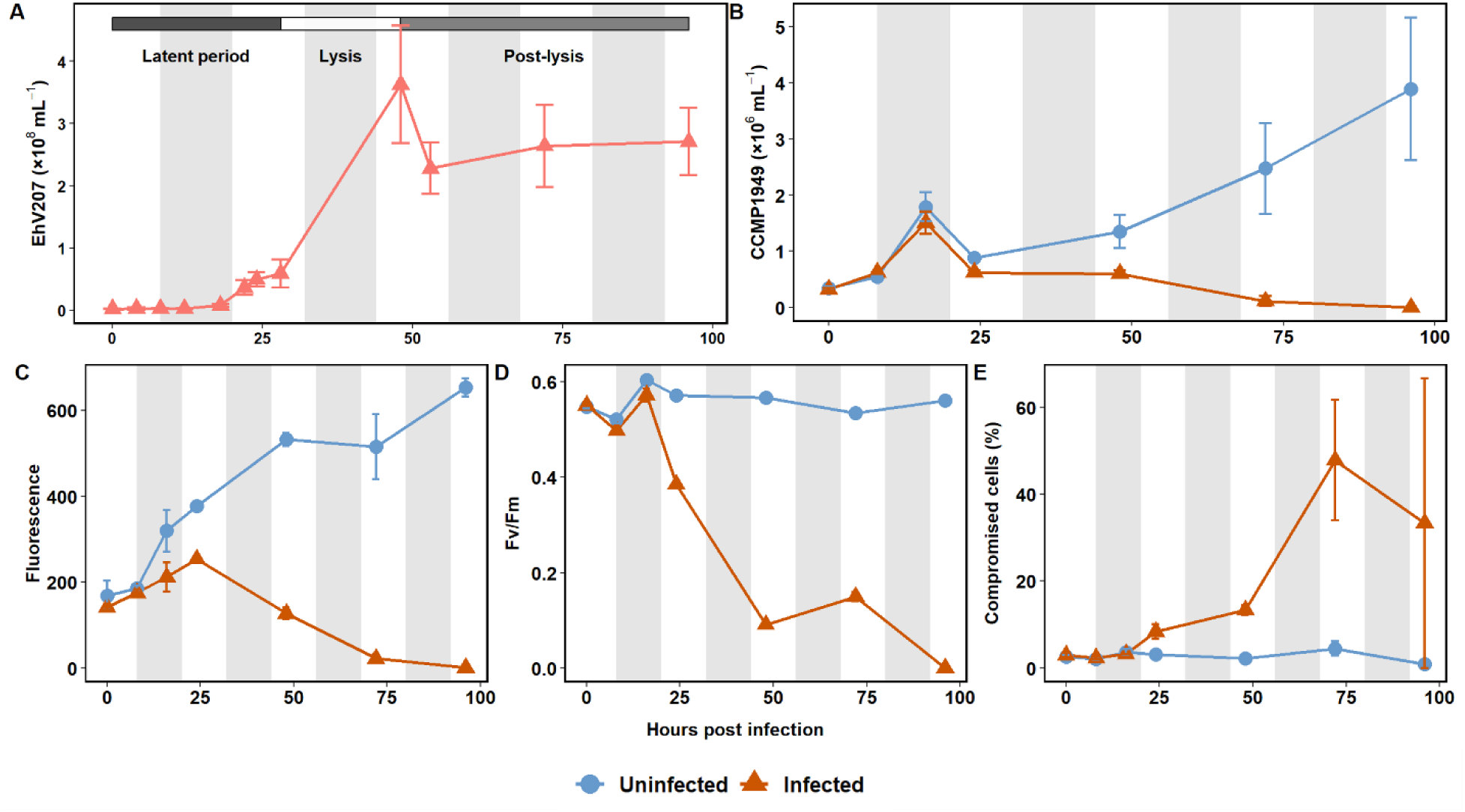
Growth of *Gephyrocapsa huxleyi* CCMP1949 in uninfected cultures and cultures infected with virus EhV207. (**A**)Viral lysis occurred between 24 to 48 hours post infection, leading to a marked increase in extracellular virus concentrations. (**B**) Cell densities of infected cultures decreased 3-orders of magnitude between 24 to 96 hours post infection. (**C-E**) No increase in compromised cell membranes or declines in fluorescence was observed during the latent period of infection. Arrows indicate when sacrificial VOC measurements were made; note that the growth data between VOC measurements is not continuous and was obtained on separate sacrificial replicates. Grey shaded areas represent dark periods.

### Viral infection increases total VOC release in *G. huxleyi*

VOCs were released rapidly as one large “pulse” during the late infection stages 24-48 h post infection, coinciding with cell lysis (Figure 1). Lytic viruses increased the total VOC pool by a maximum of approximately 6-fold within 48 hours post infection (986±124 ppbv in uninfected cultures to 6003±167 ppbv in infected cultures) (Figure 2). This represents an increase in VOC concentrations of nearly an order of magnitude compared to the time of infection (667±25 ppbv in infected cultures). Following lysis, the VOC pool declined within two days until it returned to pre-infection levels, matching VOCs of uninfected cultures. The open experimental system used for this experiment allows dissolved VOCs to escape, which aligns with our results, as total VOC concentrations in uninfected cultures remained relatively stable for most of the experiment (ranging from 753±73 to 986±124.5 ppbv). Thus, the reported VOC concentrations likely reflect production rates in equilibrium with emissions.

**Figure 2.**
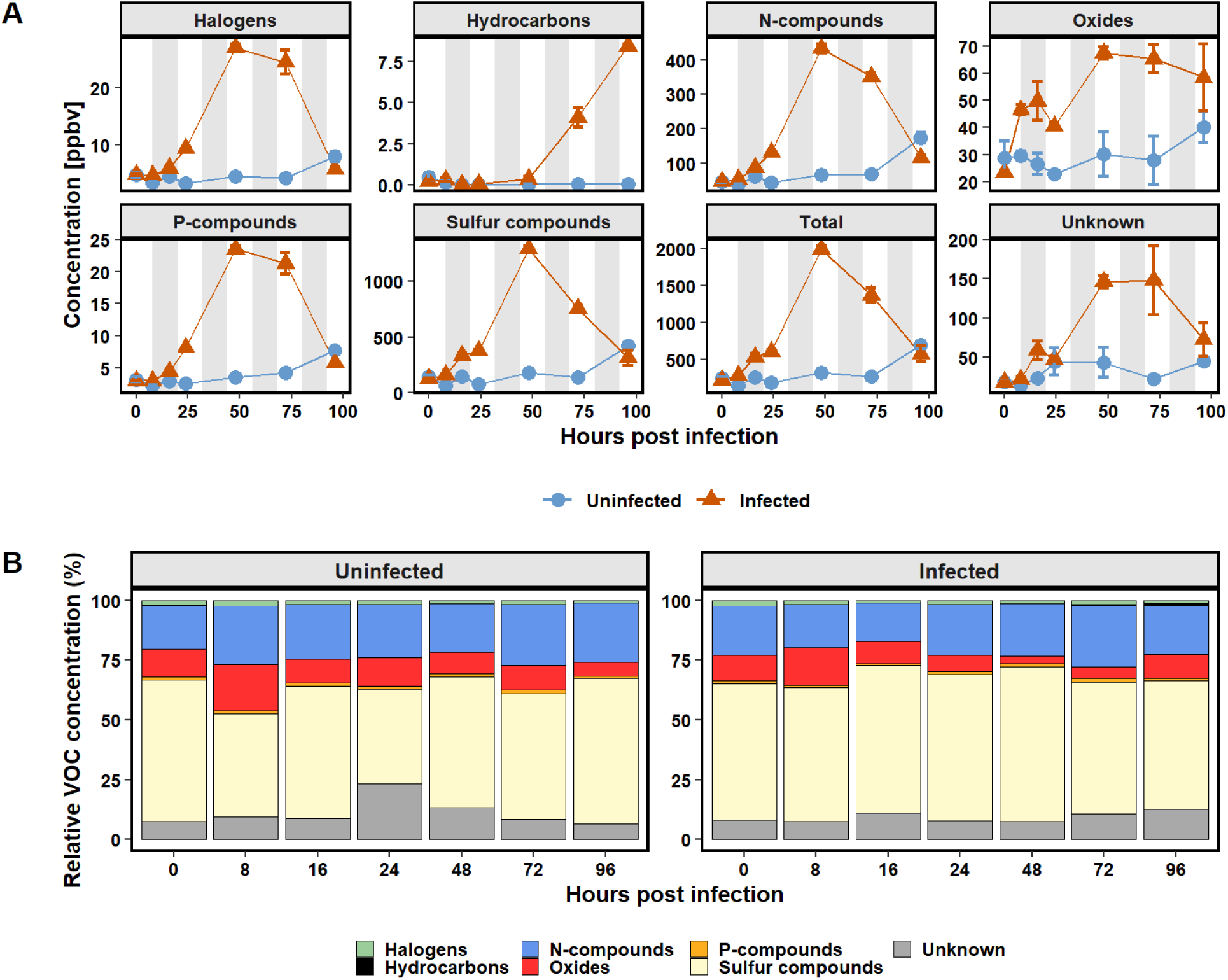
(**A**) Mean VOC concentrations (ppbv) for major groups of VOCs in infected and uninfected *Gephyrocapsa huxleyi* cultures. Error bars represent standard deviation, calculated separately for both PTR modes. (**B**) The relative proportion (%) of all VOC groups per timepoint. Relative contribution of each VOC group remained consistent throughout the experiment for both infected and uninfected cultures, despite total VOC concentrations increasing significantly in infected cultures after 24 hours. VOC release was drastically higher in virally treated cultures. Grey shaded areas represent dark periods.

The VOC pulse occurred only after viral lysis began in infected cultures; but VOCs also increased when growth rates declined in uninfected cultures after 72 hours. Halsey and Giovannoni (Halsey and Giovannoni, 2023) hypothesized that environmental perturbations could enhance the likelihood for accumulated VOCs to be emitted to the atmosphere. Nitrogen limitation in microalgae can induce VOC production (Kumari *et al*., 2020), underscoring the broader significance of VOC release during viral infection and otherwise perturbed coccolithophore populations. The implication is that resource scarcity, or the physiological stress induced by it, as well as viral predation contribute to elevated VOC release in *G. huxleyi*.

Bacteria known to have a specialized C1 metabolism are slow growing oligotrophs, such as SAR11 and OM43 (Sun *et al*., 2011; Halsey *et al*., 2012), and are thought to play an important role in biological VOC cycles(Halsey *et al*., 2017; Moore *et al*., 2020, 2022). If VOC residence times are less than 48 hours, as our findings suggest, C1 specialists would be too slow growing to take full advantage of short-lived VOC pulses. Thus, many VOCs accumulated during viral lysis could be removed from the microbial food web by flux to the atmosphere. Atmosphere-ocean surface fluxes for many VOCs such as isoprene, acetone, acetaldehyde and DMS are typically a net positive, and are correlated to their photic production (Kettle and Andreae, 2000; Sinha *et al*., 2007; Dixon *et al*., 2013). Viral induced VOC production may be an important source of VOC fluxes to the atmosphere that also contributes to spatial and temporal VOC variability (Exton *et al*., 2012; Yu and Li, 2021). In contrast, steady VOC production during phytoplankton growth may be a more significant source of nutrition for heterotrophic microbes.

### Relative contributions of VOC constituent groups in G.huxleyi host-virus systems are constant throughout the infection cycle

Out of 376 unique peaks from PTR-MS data, 20 peaks with maximum concentrations higher than 1 ppbv could be matched with known VOC compounds using primarily the GLOVOCS database for comparison (Yáñez-Serrano *et al*., 2021). 13 out of 20 of these main VOCs were not previously detected in *G. huxleyi* (Table 1). Further 57 VOCs were identified with concentrations less than 1 ppbv, which we considered trace VOCs. Binning VOCs into groups based on what elements they contained showed that despite strong differences over time in the total VOC production, the relative contribution of VOC groups (N-, S-, P-, O-containing and halogenated VOCs) remained remarkably stable regardless of infection stage (Figure 2), with no differences in the percentages of VOC groups between time points (t-tests, p<0.05). However, when considering individual VOC species, 70 peaks were found to be significantly elevated in infected cultures (Figure 3) (comparing log2fold changes in concentration between the treatments; p-adj.<0.05, determined via Wald tests). This is evidence that viral infection influences at least 18.6% of the VOC biochemistry in the *G*. *huxleyi* host.

**Figure 3.**
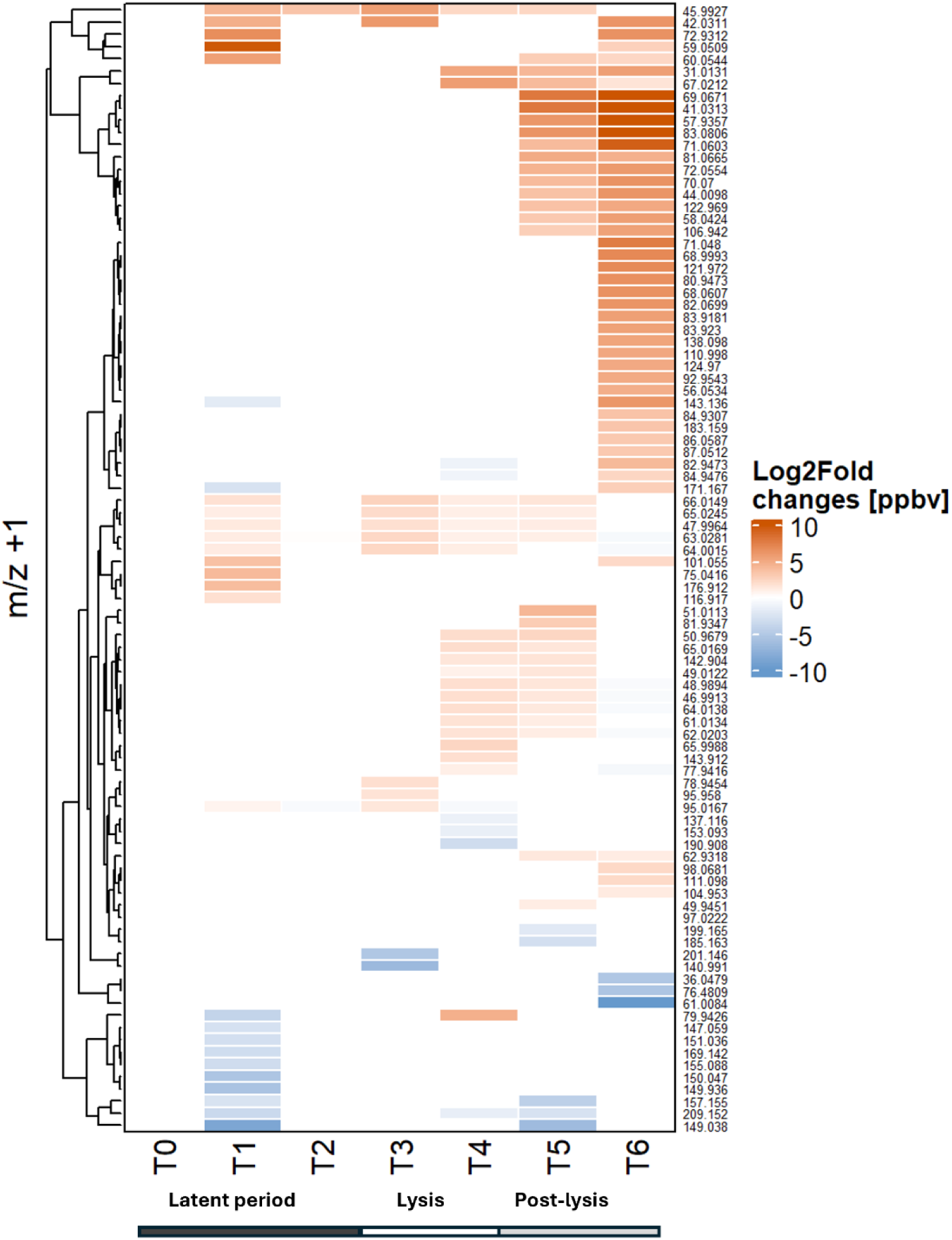
Comparison of relative changes in concentration per VOC species between EhV207 virus-treated *Gephyrocapsa huxleyi* CCMP1949 and no-virus control cultures over the course of the infection (Log2fold difference of estimated ppbv per m/z +1 value). Masses where the difference was not significant (p-adj > 0.05) are not shown. Masses in each row are shown as m/z+1. Different sets of VOCs were observed to be different between the treatments during the incubation period, lysis, and post lysis time frame (T_0_= 0 hours, T_1_= 8 hours, T_2_ = 16 hours, T_3_ = 24 hours, T_4_ = 48 hours, T_5_ = 72 hours, T_6_ = 96 hours). The largest dissimilarity was observed after lysis had concluded, however changes in VOC profiles occurred starting 8 hours after infection, suggesting viruses alter VOC release of infected *G. huxleyi* prior to lysis.

**Table 1.**
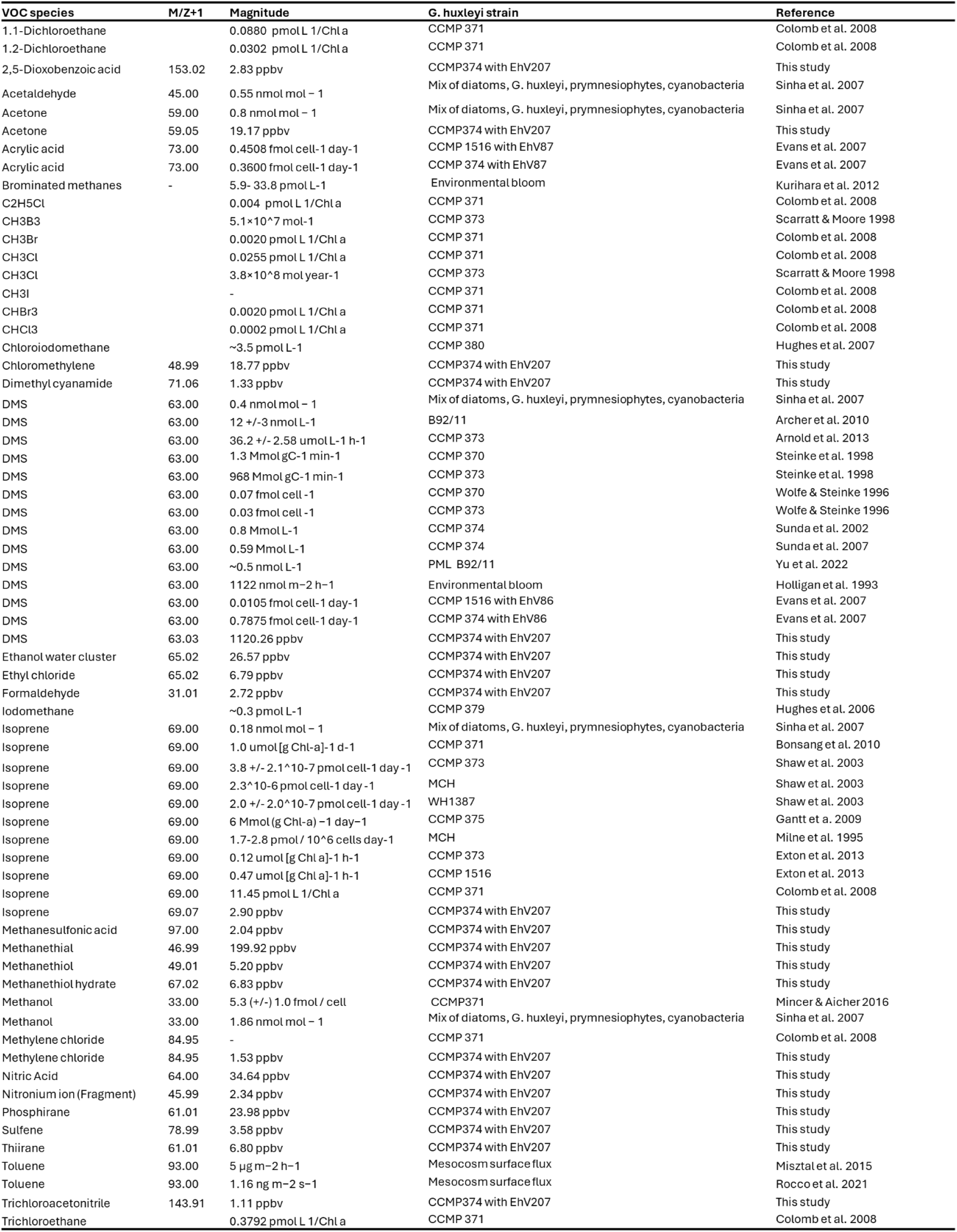
VOCs associated with *G. huxleyi* as reported in the literature and this study. We identified 79 unique VOCs in G. huxleyi cultures. 20 of those VOCs have been reported before. VOCs with less than 1 pbbv were omitted from the table for clarity.

The relative composition of VOC groups did not shift in the early infection stages (Figure 2), when the virus would be expected to hijack cell metabolism (Warwick-Dugdale *et al*., 2019). Like all viruses, lytic EhVs have been shown to rapidly and selectively shut down host transcription, altering the cellular machinery (Rosenwasser *et al*., 2019; Ku *et al*., 2020), albeit with a particular emphasis on rewiring lipid metabolism by depleting host-specific sterols (Rosenwasser *et al*., 2014). EhVs can arrest nutrient uptake of *G. huxleyi* cultures within 24 hours of infection, leading to an enrichment in C and P relative to N in the POC fraction due to lipid accumulation (Dikstein *et al*., 2024). However, the same trend could not be observed in VOCs. To further investigate the VOC compositional differences between treatments and timepoints, we used a PCoA ordination to visualize samples (Figure 4). PERMANOVA analysis found statistically significant differences in VOC composition between treatment and infection stage (p<0.01), showing that infection stage explained the largest magnitude of differences between samples (R2=0.47), whereas the model was a poor fit for treatment (R2=0.09). For infected cells, the PCoA suggests a stronger separation in VOC composition between time points than the uninfected controls. Infected *G. huxleyi* produced the same VOCs present in uninfected cells, suggesting that EhV207 does not induce or shut down host metabolic pathways involved in VOCs production during infection. Instead, viral infection increases the rates of VOC production. The largest dissimilarity in VOC concentration was observed after lysis had concluded, however changes in VOC profiles occurred starting 8 hours after infection, suggesting viruses alter VOC release of infected *G. huxleyi* prior to lysis (Figure 3). The implication is that EhV207 uses intracellular resources and host metabolic pathways, that may be significantly tied to the VOC biochemical pathways. The fact that VOC concentrations increased in a strong pulse during late-infection could be explained if (1) VOC production associated with lysis is a consequence of the virus accelerating metabolite fluxes through *G. huxleyi* biochemical pathways during the late infection stages, or (2) viral infection frees pre-existing cell bound VOCs within the cell when they lyse the membrane.

**Figure 4.**
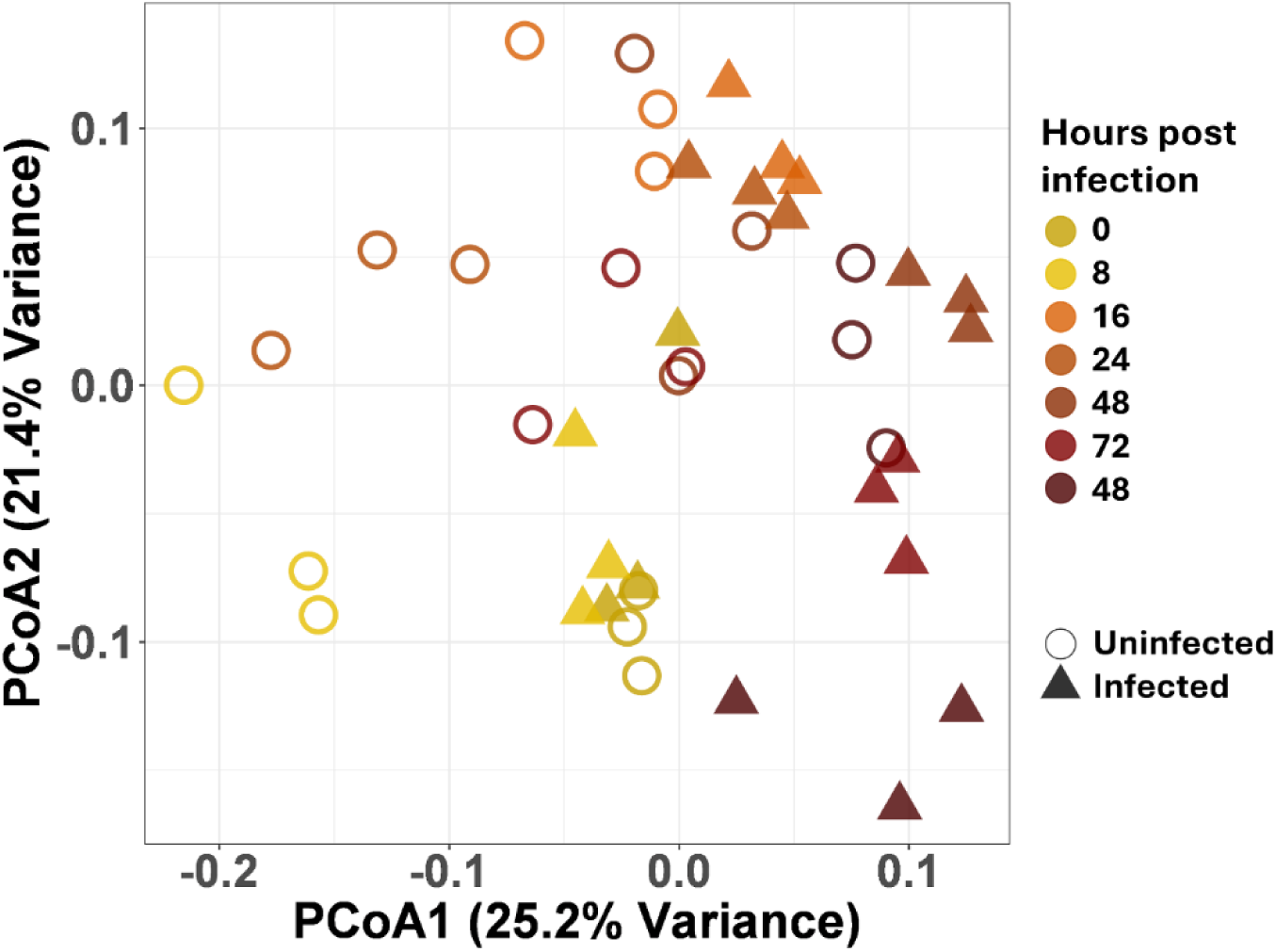
Principal component analysis shows differences when comparing all produced VOCs between uninfected and infected cells during the lysis (24-48 hours post infection) and post-lysis period (48-96 hours post infection).

It has been hypothesized that some VOCs are intermediates of cell metabolism that diffuse through membranes because of their hydrophobicity and low molecular weight (Halsey *et al*., 2017; Halsey and Giovannoni, 2023). We matched CHEBI, PubChem and KEGG compound ID’s for the VOCs that could be identify from our PTR-MS data, and mapped them against KEGG pathways for *G huxleyi*. Thirty-six VOCs could be matched with KEGG compounds, 16 of those were present in *G. huxleyi* metabolic maps (Supplementary Table 1). Therefore, there are 20 KEGG compounds that have not previously been associated with the *G. huxleyi* reference genomic maps; however, no novel pathways were found. When comparing the KEGG compounds with VOC concentrations, 70% of VOCs were related to sulfur metabolism, with the second most abundant (2%) being related to nitrogen metabolism, and remaining pathways related to less than 1% of the combined VOC concentrations. However, we could not match many abundant VOCs with KEGG compounds, e.g. CH_3_NO_2_ (m/z 62.02), as the conformation of this exact compound cannot be determined with PTR-MS. KEGG analysis was therefore congruent with the VOC release patterns, but neither offers a clear understanding of the metabolic origin and purpose of virally released VOCs.

### Halogenated VOCs from infected and uninfected *G. huxleyi*

We identified 17 halogenated VOCs in both infected and uninfected *G. huxleyi* cultures (Figure 5) with a total concentration of 14.4±0.5 ppbv at the time of the initial infection. Concentrations increased to a total of 81.3±1.9 ppbv 48 hours post infection, compared to the maximum of 23.8±2.7 ppbv after 96 hours in uninfected cells. Only nine out of the 17 halogenated compounds increased in the late infection stage (Figure 5). Previous reports showed metabolites that are both chlorinated and iodinated, such as chloroiodomethane, are produced by infected *G. huxleyi* (Fuse *et al*., 2003; Hughes *et al*., 2006). Kuhlisch et al. (Kuhlisch *et al*., 2021) suggest that halogenation of single compounds with both chlorine and iodine is a hallmark of viral *G. huxleyi* DOM, because halogens increased in exometabolites containing different combinations of two or three halogen atoms (chlorine, bromine and iodine) in a post *G. huxleyi* bloom. We identified trace amounts of eight halogenated VOCs (less than 1 ppbv) with two or more halogen atoms, but we did not identify compounds with multiple iodine or bromine atoms, although we cannot rule out they were present in the unidentified fraction. However, we identified five fluorinated compounds; and although we are unaware of any other reports of fluorinated VOCs associated with *G. huxleyi* blooms, fluorine containing metabolites have been attributed to phytoplankton communities, e.g. trichlorofluoromethane and dichloromethane production in the Indian ocean (Ooki and Yokouchi 2011), and trichlorofluoromethane, among other halocarbons, in the Yangtze River Estuary (Zou *et al*., 2022).

**Figure 5.**
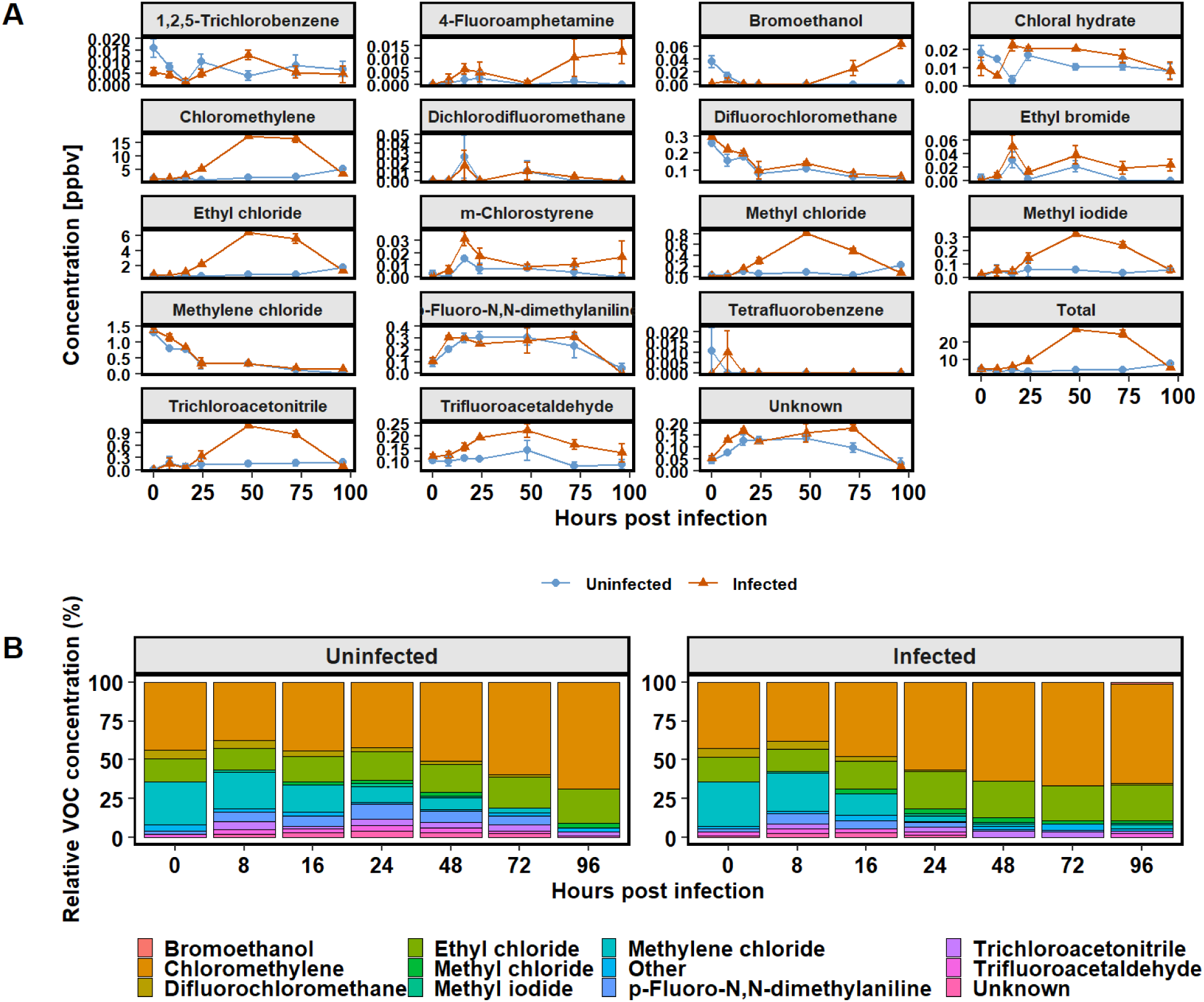
Estimated concentration of halogenated VOCs in infected and uninfected *G. huxleyi* cultures. (**A**) Mean concentration (ppbv) of all detected halogens over the course of the experiments, with error bars indicating the standard error. (**B**) Relative contribution of compounds per timepoint, trace compounds (less than 1 % contribution to total) were grouped as “Other”. Compounds binned into “Unknown” were due to structural ambiguity of identical molecular masses, which did not allow for identifying exact molecular configuration of those masses. Chloromethylene and ethyl chloride constituted more than half of all detected halogens throughout. Despite a strong pulse of halogens released via viral lysis, the proportions of halogens between treatments did not change substantially.

Halogenated VOCs constituted approximately 2% of all VOCs in both treatments; similar to the pattern of unhalogenated VOCs, halogenated VOCs rose sharply in the late infection stages, although the ratio of unhalogenated to halogenated compounds did not change with infection status. As none of the observed halogenated VOCs were unique to infected cultures, there is no evidence to suggest that EhVs are able to imbue their host with that metabolic capability. The first halogenase reported in a marine virus was found recently in a cyanophage (Gkotsi *et al*., 2019), and no halogenases have been reported from EhV. Mono-methylhalides are produced by phytoplankton (Scarratt and Moore, 1998; Sœmundsdóttir and Matrai, 1998) using halide ion methyltranferase and S-adenosine methionine as the methyl donor (Ohsawa *et al*., 2001; Itoh *et al*., 2009). Halogenated metabolites associated with EhVs may be a result of enhanced haloperoxidase activity in *G. huxleyi* cells (Kuhlisch *et al*., 2021) as a strategy to scavenge surplus reactive oxygen species during viral infection. These extracellular haloperoxidases catalyze oxidation of a halide anion by peroxide, yielding electron-deficient halide ions that halogenates DOM, yielding complex organohalides

(Paul and Pohnert, 2011). Haloperoxidase activity consumes peroxide that can accumulate in phytoplankton under high light conditions (Pedersen *et al*., 1996). A role in grazing defense has also been suggested (Paul *et al*., 2006), thus organohalide production may protect cells against stress. However, it is unclear whether organohalides could aid cells against viral predation. Furthermore, there was no increase in halogenated VOCs during the latent period, only during the late infection stage when the virus would have likely overcome any viral defenses. These results indicate that halogenated VOCs probably do not play a role in cell defense against viral infection or stress in this virus-host system. Alternatively, we hypothesize that EhVs shuttle halogenated compounds from the POM to the DOM pool via cell lysis, rather than increasing halogenation of DOM directly.

### Coccolithophores release diverse sulfur VOCs

Coccolithophores produce diverse and abundant sulfur VOCs, such as dimethylsulfoniopropionate (DMSP), dimethylsulfide (DMS) and sulfonate 2,3-dihydroxypropane-1-sulfonate DHPS (Evans *et al*., 2007; Durham *et al*., 2019). Our analysis identified 18 sulfur-containing VOCs, two of which were classified as “unknown” due to multiple possible molecular arrangements (Figure 6). Sulfur-containing compounds were the most abundant in both infected and uninfected *G. huxleyi*, with up to three times the amount compared to other VOC groups. Sulfur compounds in infected cultures increased from 125.8±4 ppbv to 1293±30 ppbv within the first 48 hours and decreased to 312±66 ppbv after 96 hours. Control cultures remained steady from 0 hours (147.2±12 ppbv) to 72 hours (141±3 ppbv) but increased to 426±21 ppbv by 96 hours. Sulfur VOCs were more abundant after 16 hours, prior to increases in viral abundance, whereas halogen-, phosphorus-, and nitrogen-containing VOCs increased coincident with virus release. Therefore, some sulfur VOCs were released from infected cells prior to lysis. However, as the ratios of sulfur VOCs to total VOCs remain similar throughout the infection cycle, and our results indicate limited change in metabolic capacities, suggesting that either these compounds have different biophysical properties that allow them to diffuse out of the cell more easily, or that a potentially much larger intracellular pool of sulfur VOCs increases the osmotic gradient increasing diffusion through the cell membrane. The presence of large intracellular sulfur VOC pools would support our hypothesis that viral infection frees pre-existing cell bound VOCs within the cell when they lyse the membrane.

**Figure 6.**
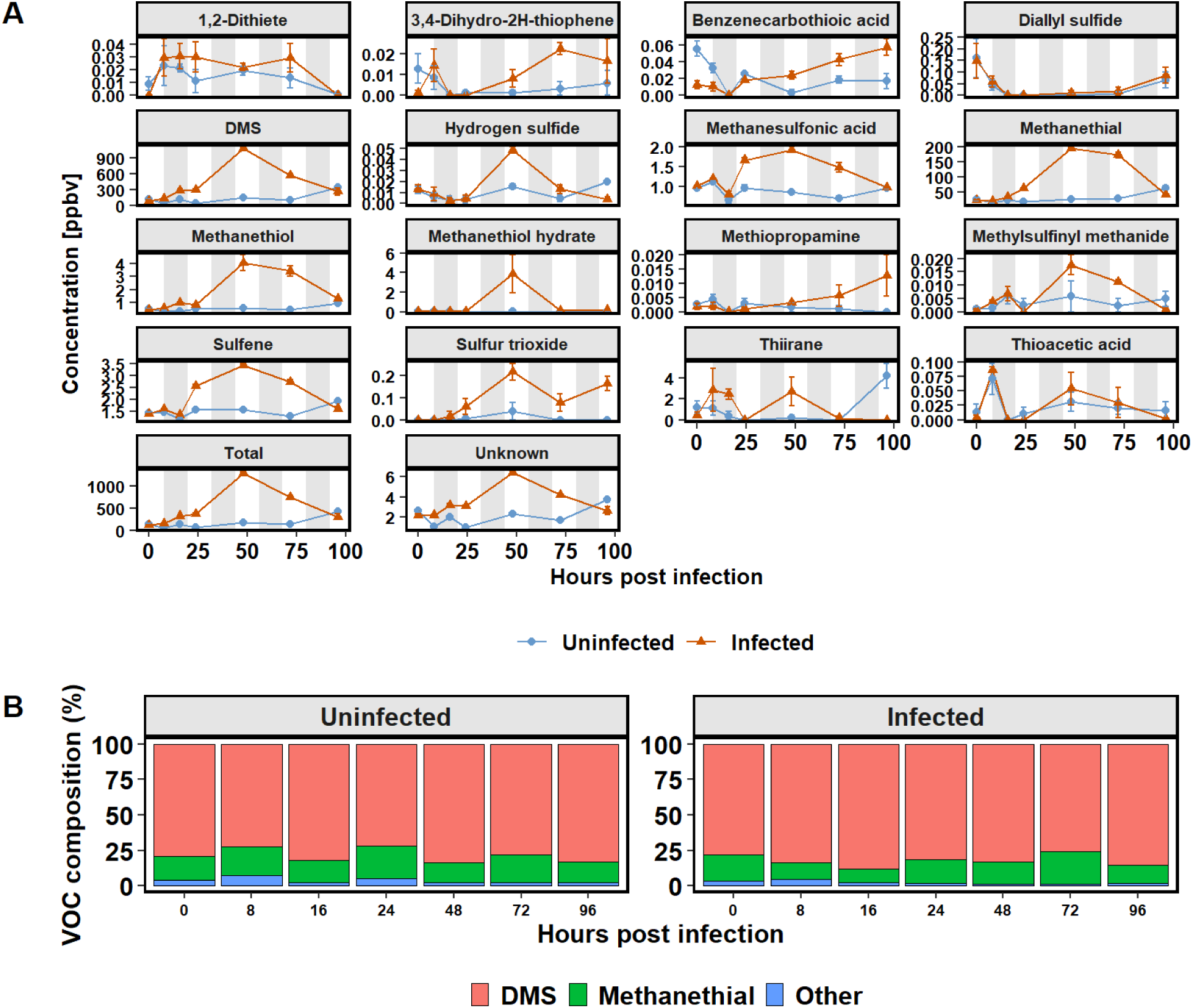
Estimated concentration of sulfur VOCs in infected and uninfected *G. huxleyi* cultures. (**A**) Mean concentration (ppbv) of all detected sulfur compounds over the course of the experiments, with error bars indicating the standard error. (**B**) Relative contribution of compounds per timepoint, trace compounds (less than 1 % contribution to total) were grouped as “Other”. Unlike with other VOC groups, a single compound (dimethyl sulfide (DMS)) was clearly more predominant in all cultures, marking it as the most important VOC in *G. huxleyi* (by volume).

DMSP, a precursor to dimethyl sulfide (DMS), is an important and abundant intracellular osmolyte in marine algae (Steinke *et al*., 1998; Stefels *et al*., 2007). DMS accounted for up to 85% of sulfur-containing VOCs in infected *G. huxleyi*, and 82.8% in uninfected cultures (Figure 6), which was roughly half of all VOCs regardless of infection stage or status (Figure 1). The magnitude and distribution of DMS release by viral lysis in these experiments was comparable to previous reports (Evans, Malin, Wilson, *et al*., 2006; Evans *et al*., 2007); however, the ratio of DMS to other VOCs did not change as a result of infection. It has been well established that grazing and viral infection on DMSP-producing algae, like *G. huxleyi*, induces DMS and acrylic acid via DMSP cleavage, possibly to deter grazers and reduce viral titers (Malin *et al*., 1998; Evans, Malin, Wilson, *et al*., 2006; Evans *et al*., 2007; Todd *et al*., 2010; Curson *et al*., 2018; Simó *et al*., 2018). Although DMSP lyase activity as lower in infected cells compared to grazed cells (Evans *et al*., 2007), total DMS production was thought to be higher in declining *G. huxleyi* blooms (Le Gland *et al*., 2024). If the spike of DMS observed here, and in the literature, was the result of virally manipulated DMSP cleavage, as is often suggested, the ratio of DMS compared to other VOCs should increase in infected cultures compared to controls. In contrast, the ca. 80% DMS-sulfur VOC to total VOC ratio did not change over the course of the experiment and remained similar to ocean observations, where DMS constitutes 70% of sulfur emissions (Carpenter *et al*., 2012).

In addition to DMS, DMSP cleavage can also free methanethiol (MT) (Howard *et al*., 2008; Reisch *et al*., 2011), which is typically the second most abundant organosulfur compound in the marine environment (Halsey *et al*., 2024). We observed small increases in MT and MT hydrate similar to DMS (1.2±0.05 ppbv MT 16 hours post infection). Specifically, the majority of MT production correlated to viral particle release (4.0±0.6 ppbv and 3.8±1.9). Less than 1 ppbv MT was observed in uninfected control cultures, suggesting viral infection induces MT in *G. huxleyi*. MT has been reported from PTR-MS head-space analysis of axenic *Thalassiosira pseudonana*, which is a producer of sulfur VOCs, comparable to *G. huxleyi (Kameyama et al., 2011)*. Unlike DMS, Kameyama et al. (Kameyama *et al*., 2011) reported a strong correlation of MT with the photic period. We are not aware of reports of MT production in *G. huxleyi,* or in association with algal viruses, but MT production has been detected during a coccolithophore-dominated phytoplankton bloom in the Atlantic Ocean (Davie-Martin *et al*., 2020), suggesting *G. huxleyi* blooms may be a source of biological MT (Lawson *et al*., 2020). Lawson et al., (Lawson *et al*., 2020) provided evidence for positive surface flux of MT (3.5 to 5.8 μmol m⁻² d⁻¹) in the Pacific Ocean, which was 14% to 24% of the DMS flux measured (MT / (MT + DMS)). Marine heterotrophic communities produce MT by catabolizing DMSP into DMS and MT (Sun *et al*., 2016), therefore MT is often attributed to bacterial DMSP lyase activity. *G. huxleyi* also has DMSP cleaving enzymes (Alcolombri *et al*., 2015), although this pathway is not known to release MT. Our results suggest that viral predation frees MT, however the biochemical mechanism remains unknown. Even though MT concentrations were low compared to DMS, MT metabolic pathways are undescribed in coccolithophores and our observations indicate they may be contributors to MT cycling.

Surprisingly, we observed thioformaldehyde (methanethial, CH_2_S) as the second most abundant sulfur compound released (max. 195.6±3.8 in infected cultures after 48 hours). Microbial production of thioformaldehyde from potato and prawn microbiomes have been reported (Chinivasagam *et al*., 1998; Vita *et al*., 2015), but this is the first reports to link it to phytoplankton. Thermal decomposition of DMS can form thioformaldehyde (Johnson *et al*., 1971), which may explain some of the observed thioformaldehyde, as the PTR-MS reaction chamber is at ∼80°C. The ratio of thioformaldehyde to DMS increased strongly with higher DMS concentrations, from 2.6±0.3 (100 nM DMS) to 72±0.8 (5 μM DMS) ppbv DMS per unit thioformaldehyde in DMS standards (Supplementary Figure 2), suggesting that higher DMS concentration increase rates of decomposition. If thermal decomposition of DMS was the source of thioformaldehyde in *G. huxleyi* cultures, the concentration of DMS would be correlated to thioformaldehyde. However, the thioformaldehyde signal was not consistently proportional to DMS concentrations in *G. huxleyi* cultures, ranging from 3.3±0.3 to 9.0±1.4 ppbv DMS per unit thioformaldehyde. Therefore, it seems likely that at least part of the observed thioformaldehyde is biologically produced by *G. huxleyi*.

### Potential nitromethane production in G. huxleyi

We observed 19 nitrogen-containing VOCs with unidentified structures, mostly in trace amounts (<1 ppbv) (Supplementary Figure 3). Over 90% of the unidentified compounds were CH_3_NO_2_ (m/z+1 62.02), which could be nitromethane or methyl nitrite depending on its structure. Either compound would be surprising, as nitromethane is a biofuel and typically produced by combining propane with nitric acid, while methyl nitrate results from the condensation of nitric acid and methanol. Neither are known to be produced by *G. huxleyi*. Additionally, a peak at m/z 43.01 was observed, which could represent n-propane fragments and was relatively abundant (36.4±7.2 ppbv 16 hours after infection). Nitric acid was the second most abundant nitrogen compound, and its abundance hints at a mechanism for CH_3_NO_2_ production in *G. huxleyi*.

Marine alkyl nitrates, including methyl nitrate, are natural sources of atmospheric nitrogen oxide and contribute to tropospheric ozone formation (Chuck *et al*., 2002; Moore and Blough, 2002; Neu *et al*., 2008). The release of nitrogen-containing VOCs, such as methylamines and acetonitrile, has been linked to DOM photolysis and bacterial metabolism (Moore and Blough, 2002). Their production may occur via photochemical reactions between alkyl groups and NO (Dahl *et al*., 2003). Green microalgae can reduce nitric oxide (NO) to nitrous oxide (N2O), an important greenhouse gas. Schieler et al. (Schieler *et al*., 2019) showed that *G. huxleyi* viruses produce NO as a marker of early-stage infection, which may have antioxidant properties that facilitate viral replication by preventing reactive oxygen species buildup. During phage infections, nitrate reductase activity remains constant in cyanobacteria, supporting high protein demands for viral reproduction (Bisen, 1986). Though alkyl nitrates are known, we could not find an explanation for CH_3_NO_2_, but it reveals an exciting possibility for biotechnological applications.

## Conclusion

This study demonstrates the profound impact of viral infection on VOC emissions in *G. huxleyi*. Infection by EhV207 increased VOC production nearly six-fold. We identified 13 VOCs with concentrations higher than 1 ppbv not previously detected in *G. huxleyi* and 79 VOCs in total were significantly impacted by viral infection, including 15 sulfur, 22 nitrogen, 2 phosphorus, 19 oxygen and 17 halogen-containing compounds, with sulfur-containing VOCs, such as DMS, dominating the profiles. Furthermore, our results suggest that infection stages do not alter the relative contribution of different VOC groups, despite a sharp increase in concentration of individual compounds, correlating to virus release 24–48 hours post-infection. The lack of unique viral VOCs, coupled with the consistent ratios, indicates that EhV207 primarily accelerates existing metabolic processes and facilitates the release of pre-existing intracellular VOCs rather than inducing novel biochemical pathways. These findings underscore the ecological significance of viral infections in marine systems, emphasizing the potential of viral lysis in enhancing VOC flux to the atmosphere and suggest that viral-induced VOC pulses contribute to VOC variability and biogeochemical cycles. Overall, the study provides new insights into the interactions between viral infections, phytoplankton metabolism, and VOC emissions, highlighting the broader implications for marine-atmosphere interactions and VOC cycling. Further investigations into the temporal dynamics of VOC release and their atmospheric fate will enhance our understanding of the role of viral lysis in shaping marine ecosystem processes.

## Materials and Methods

### E. huxleyi cell and virus culture conditions

Cultures of *G. huxleyi* CCMP374 (https://ncma.bigelow.org/CCMP1949) were used for all laboratory experiments. CCMP374 (synonym CCMP1492) is a naked strain originally isolated in the Gulf of Maine, which is frequently used as a model organism for G*. huxleyi* due to its high sensitivity to viral infection (Bidle and Kwityn, 2012). Regular stock cultures were maintained at 16°C without rotation in 250 mL PC culture flasks using the defined F/2 media (Guillard, 1975). As a light source we chose a 1:1 mix of blue and red lights at peak wavelength 420 nm and 650 nm on a 14:10 h light-dark cycles (∼150 μE m^-2^ s^-1^) as suggested for optimum growth in *G. huxleyi* (Granata *et al*., 2019). For PTR-MS experiments, a singular 100 mL starter culture was used to inoculate 110 mL F/2 media in 250 mL PC flasks to 1×10^5^ cellsmL^-1^. Cultures were maintained in the same conditions as the stock cultures until they reached 5×10^5^ cellsmL^-1^ to ensure cells were acclimated and in the same physiological state before starting the experiment.

Cultures of the model *G. huxleyi* virus EhV207 were generously provided by the Knowles lab at UCLA. EhV207 infections result in rapid lysis in host cultures and are considered a highly virulent virus for CCMP1949 (Nissimov *et al*., 2016; Knowles *et al*., 2020). EhV207 was maintained by addition of virus to exponentially growing host culture with cell density of 5×10^5^ cells mL^-1^ at a 10:1 MOI. Cultures typically visibly cleared within 2-3 days after infection, after which cells were counted using flow cytometry and compared to uninfected parallel control cultures. Cell debris was removed with a 0.45 µm syringe filter (Sterivex filters Millipore Sigma, Burlington, MA, United States). Fresh lysate was prepared in the same fashion before each experiment, stored in the dark at 4 °C, and used within one week.

### *E. huxleyi* cell counts and staining

Cell density of *G. huxleyi* was measured on cell autofluorescence using a Guava Technologies flow cytometer (Millipore; Billerica, MA). The acquisition settings were established prior to the experiment using non-infected cells: Events were gated with log_10_-transformed red-fluorescence (488/692 nm excitation/emission) plotted against log_10_-transformed side scatter inside the Incyte software. To count samples, 200 µL culture were transferred to 96-well flat bottom plates, without fixation and measured at room temperature for 30 seconds per sample. Events counted within the established gates were calculated to cell densities using flow rates per minute. Dead cells and cells with compromised membranes were detected and enumerated using flow cytometry and the live-dead stain SYTOX Green (ThermoFisher S7020, USA) stained cells. 5 mM SYTOX Green stock solution was added to 200 µL culture aliquots in 96 flat-bottom well plates to a final concentration of 1 µM and incubated in the dark at room temperature for 30 minutes before analyzing samples on the Guava Technologies flow cytometer. Counting gates were applied to plots of log_10_-transformed side scatter against log_10_-transformed green fluorescence (488/520 nm excitation/emission). Event counts from SYTOX Green stained cells were then compared to autofluorescence cell counts to estimate the percentage of dead or membrane compromised cells.

### Viral abundance

Viruses were enumerated using a Becton-Dickinson FACSCelestra flow cytometer equipped with a violet (405nm; 50 mW), blue (488nm; 20mW), and red (640nm; 40mW) laser configuration according to Marie et al. (Marie *et al*., 1999), with modifications by Mojica et. al. (Mojica and Brussaard, 2014). Samples were fixed with 25% glutaraldehyde (EM-grade, Sigma-Aldrich) at a final concentration of 0.5% for 15–30 min at 4 °C, flash frozen, and stored at − 80 °C until analyzed. Thawed samples were diluted using TE buffer (pH 8.2, 10 mM Tris-HCL, 1 mM EDTA; Sigma-Aldrich) to achieve an optimum event rate of 200-800 events/second. Virus samples were stained by heating in the dark at 80 °C for 10 min in the presence of SYBR Green I at a final concentration of 0.5 × 10−4 of the commercial stock concentration. Stained samples were discriminated and enumerated based on their 488 nm excited right-angle light scatter (SSC) and green fluorescence (530 nm). The list mode files obtained from FC were analyzed using the software package FCS Express (v7).

### Detection of Volatile Organic Compounds

For a qualitative estimate of Volatile Organic Compound (VOCs) 100-mL samples were transferred to custom-made 200 mL polycarbonate dynamic stripping chambers with sintered glass frits (2-2.5 µm) which were previously designed to measure VOCs in cell cultures (Halsey *et al*., 2017; Davie-Martin *et al*., 2020). The VOCs were stripped from the sample inside the stripping chamber using 50 sccm (Sierra Instruments, Monterey, CA, United States) of carrier gas (Zero grade air) which was flown through a hydrocarbon trap and then piped through the glass frits inside the stripping chamber to establish a uniform pattern of fine bubbles. The VOC infused carrier gas was vented into a Proton Transfer Reaction time of flight mass spectrometer (PTR-TOF-1000 MS, IONICON analytik GmbH, Innsbruck, Austria) using teflon tubing. Two different settings were used for both sets of samples to capture different types of VOCs. The first set used hydronium (H_3_O^+^) ions created from the PTR-MS internal water reservoir for a soft ionization event to VOCs having higher proton affinities than 691 kJ/mol. Data in the mass range of 18 to 363 a.m.u. was collected at the VOC molecular masses plus one (*m*/*z* +1) over 60 cycles at 5000 ms (approximately 5 minutes). Drift tube conditions were 2.1 mbar, 80 °C, 500 V with an E/N value of 126 Td.

### VOC data analysis

PTRwid (Holzinger, 2015) was used to create a table of integrated m/z +1 peak signals from the raw PTR-MS peak data that incorporated a correction for overlapping peaks. This was used to create a list of unified masses of all m/z +1 values for both sets of PTR-MS assays. The raw data was manually calibrated within the PTR-MS Viewer software (v.3.4.4.) using the three-point method. Briefly, for each sample, internal standards at m/z 29.99, 203. 94 and 330.85 were used to calibrate all other peaks (Gauss fit, averaging Spectra =3, search range (m/z 0.2)). Peaks were then adjusted with the autofit function using the unified mass list as peak table. Known contaminants and internal standards were removed from the list for downstream analysis (Supplementary Tables 2, 3). Data collected from the first 2.5 minutes (30 cycles) was removed from the dataset to account for lab air contamination introduced by loading the samples into the stripping chambers. Data collection cutoff was set at cycle 60 (5 minutes). For a putative peak assignment of m/z +1 values, we used the GLOVOCs database for PTR-MS data (Yáñez-Serrano *et al*., 2021). The processed data was then imported into the R studio statistical software version 4.2.2 (Boston, MA, United States) for analysis. Differences in estimated concentrations at m/z +1 between the virus treated cultures and controls were calculated using the DESeq2 (Love *et al*., 2014) package. PERMONOVA and ordination analysis were run using the R vegan package (https://vegandevs.github.io/vegan/). Figures were prepared using the ggplot2 (Wickham, 2016) and complexheatmap packages (Gu *et al*., 2016; Gu, 2022).

## Data Availability

All codes and VOC raw data will be made available in form of h5 files in the next version scheduled for release in March 2025.

## Acknowledgements

We thank the Simons Foundation for funding H.H.B. through a postdoctoral fellowship, and S.J.G. through the BIOS-SCOPE project. Virus counting was conducted by K.D.A.M. at USM. We thank Ben Knowles at UCLA for providing EhV207 cultures, and the NCMA at Bigelow Laboratory for providing *G. huxleyi* cultures.

## Competing interests

The authors declare no conflict of interests.

## Supplementary Information

**Supplementary Figure 1.**
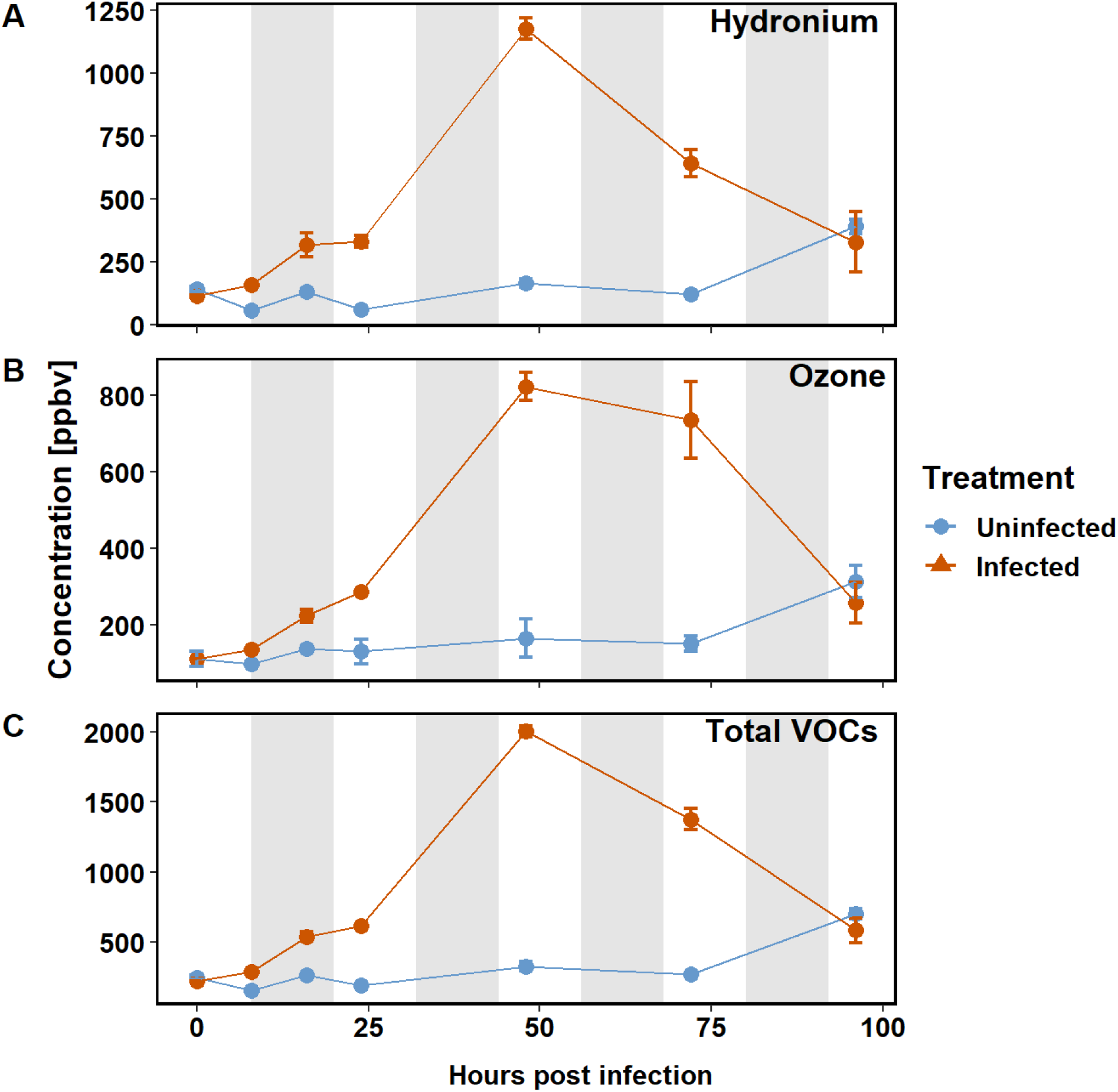
Total sum of estimated VOC concentration by ion source. (**A**) VOCs measured in the H3O+ mode, using hydronium as primary ion source; (**B**) VOCs measured in the O2+ mode, using ozone as primary ion source; (**C**) Sum of all VOC concentration estimates measure with hydronium and ozone as ion source. Error bars indicate standard deviation.

**Supplementary Table 1.** Table of KEGG compounds corresponding to VOCs. The compounds are grouped into pathways known for *G. huxleyi*, single compounds may be present in multiple pathways. Compounds which could not be assigned to a pathway are binned into “Unknown”.

| Metabolic Pathway | VOCs (ppbv) in healthy cells | VOCs (ppbv) in infected cells | Ratio | Total % of VOCs in infected cells | Total % of VOCs in healthy cells |
| --- | --- | --- | --- | --- | --- |
| Sulfur metabolism | 377.80 | 1132.82 | 3.00 | 69.25 | 68.10 |
| Unknown | 143.37 | 395.02 | 2.76 | 24.15 | 25.84 |
| ABC transporters | 10.45 | 34.64 | 3.32 | 2.12 | 1.88 |
| Nitrogen metabolism | 10.45 | 34.64 | 3.32 | 2.12 | 1.88 |
| Butanoate metabolism | 9.16 | 19.17 | 2.09 | 1.17 | 1.65 |
| Cysteine and methionine metabolism | 2.16 | 10.51 | 4.87 | 0.64 | 0.39 |
| Biosynthesis of secondary metabolites | 0.38 | 2.94 | 7.68 | 0.18 | 0.07 |
| Terpenoid backbone biosynthesis | 0.32 | 2.90 | 9.08 | 0.18 | 0.06 |
| Carbon metabolism | 0.24 | 2.83 | 12.02 | 0.17 | 0.04 |
| Nucleotide metabolism | 0.07 | 0.09 | 1.33 | 0.01 | 0.01 |
| Pantothenate and CoA biosynthesis | 0.07 | 0.09 | 1.33 | 0.01 | 0.01 |
| Pyrimidine metabolism | 0.07 | 0.09 | 1.33 | 0.01 | 0.01 |
| beta-Alanine metabolism | 0.07 | 0.09 | 1.33 | 0.01 | 0.01 |
| 2-Oxocarboxylic acid metabolism | 0.02 | 0.01 | 0.54 | 0.00 | 0.00 |
| Alanine | 0.02 | 0.01 | 0.54 | 0.00 | 0.00 |
| Biosynthesis of amino acids | 0.02 | 0.01 | 0.54 | 0.00 | 0.00 |
| Biosynthesis of cofactors | 0.02 | 0.01 | 0.54 | 0.00 | 0.00 |
| Carbon fixation by Calvin cycle | 0.02 | 0.01 | 0.54 | 0.00 | 0.00 |
| Citrate cycle (TCA cycle) | 0.02 | 0.01 | 0.54 | 0.00 | 0.00 |
| D-Amino acid metabolism | 0.02 | 0.01 | 0.54 | 0.00 | 0.00 |
| Glycolysis / Gluconeogenesis | 0.02 | 0.01 | 0.54 | 0.00 | 0.00 |
| Glyoxylate and dicarboxylate metabolism | 0.02 | 0.01 | 0.54 | 0.00 | 0.00 |
| Propanoate metabolism | 0.02 | 0.01 | 0.54 | 0.00 | 0.00 |
| Pyruvate metabolism | 0.02 | 0.01 | 0.54 | 0.00 | 0.00 |

**Supplementary Table 2.** Table of known lab and machine contaminants and their isotopic pattern peaks (IPP) that were removed from the dataset.

| Mass (m/z) removed | Type | Notes |
| --- | --- | --- |
| 30.997 | Contaminant | NO+ (ion source contaminant) |
| 31.988 | Contaminant | O2+ (ion source) |
| 32.992 | Contaminant | O2H+ (ion source) |
| 33.993 | Contaminant | H2O.H3O+ (hydrate cluster) |
| 37.044 | Contaminant | H2O.H317O+ (hydrate cluster) |
| 38.033 | Contaminant | 3H2O.H3O+ (hydrate cluster) |
| 39.033 | IPP | Mass 38.033 |
| 44.993 | Contaminant | CO2H+ (contaminant peak) |
| 47.022 | Contaminant | NO2+ (ion source contaminant) |
| 55.052 | Contaminant | 15NO2+ (ion source contaminant) |
| 57.036 | Contaminant | 2H2O.H3O+ (hydrate cluster) |
| 57.064 | Contaminant | 2H2O.H318O+ (hydrate cluster) |
| 58.072 | IPP | Mass 57.064 |
| 73.057 | Contaminant | 3H2O.H3O+ (hydrate cluster) |
| 74.036 | IPP | Mass 73.057 |
| 123.963 | Contaminant | Teflon rings (drift tube contaminant) |
| 203.955 | Contaminant | 1,3-diiodobenzene (internal calibrant) |
| 204.959 | Contaminant | 1,3-diiodobenzene (internal calibrant) |
| 205.956 | Contaminant | 1,3-diiodobenzene (internal calibrant) |
| 329.904 | Contaminant | 1,3-diiodobenzene (internal calibrant) |
| 330.931 | Contaminant | 1,3-diiodobenzene (internal calibrant) |
| 331.935 | Contaminant | 1,3-diiodobenzene (internal calibrant) |

**Supplementary Table 3.** Table of removed duplicate mass peaks. Certain VOCs may be detectable on a PTR-MS in both H3O+ and O2+ mode. To avoid biasing the analysis, the smaller peaks within the same range (mz+1 value +/- 0.005 of each other) were removed from the dataset.

| Peak m/z | Ion source | Class | Compound |
| --- | --- | --- | --- |
| 41.0275 | o2 | Non-methane hydrocarbon | Unknown |
| 43.0149 | h3o | Oxygenated VOCs | Unknown |
| 45.9925 | o2 | N-containing VOCs | Nitronium ion (Fragment) |
| 46.9916 | h3o | Sulfur VOCs | Methanethial |
| 47.9969 | o2 | Unknown | Unknown |
| 51.9429 | h3o | Unknown | Unknown |
| 52.9444 | o2 | Unknown | Unknown |
| 53.9419 | o2 | Unknown | Unknown |
| 55.9365 | h3o | Unknown | Unknown |
| 57.9328 | h3o | Unknown | Unknown |
| 59.9316 | o2 | Unknown | Unknown |
| 62.027 | h3o | N-containing VOCs | Unknown |
| 70.0765 | o2 | N-containing VOCs | Unknown |
| 71.9342 | o2 | Unknown | Unknown |
| 73.9369 | h3o | Unknown | Unknown |
| 75.9437 | h3o | Unknown | Unknown |
| 85.9342 | h3o | Unknown | Unknown |
| 87.914 | h3o | Unknown | Unknown |
| 89.9333 | h3o | Unknown | Unknown |
| 90.9457 | o2 | Sulfur VOCs | 1,2-Dithiete |
| 91.9485 | h3o | Unknown | Unknown |
| 93.9686 | o2 | Unknown | Unknown |
| 95.0175 | o2 | Sulfur VOCs | Unknown |
| 95.9649 | o2 | Unknown | Unknown |
| 101.935 | h3o | Unknown | Unknown |
| 102.939 | o2 | Unknown | Unknown |
| 103.945 | h3o | Unknown | Unknown |
| 104.95 | h3o | Unknown | Unknown |
| 105.934 | h3o | Unknown | Unknown |
| 106.94 | h3o | Unknown | Unknown |
| 107.951 | h3o | Unknown | Unknown |
| 108.965 | h3o | Halogenated VOCs | Ethyl bromide |
| 109.959 | o2 | Unknown | Unknown |
| 113.031 | h3o | N-containing VOCs | Uracil |
| 117.96 | h3o | Unknown | Unknown |
| 118.945 | o2 | Unknown | Unknown |
| 119.957 | o2 | Unknown | Unknown |
| 120.96 | h3o | Halogenated VOCs | Dichlorodifluoromethane |
| 122.967 | o2 | Unknown | Unknown |
| 125.968 | h3o | Unknown | Unknown |
| 126.897 | h3o | Unknown | Unknown |
| 134.96 | h3o | Unknown | Unknown |
| 143.914 | h3o | Halogenated VOCs | Trichloroacetonitrile |
| 145.887 | o2 | Unknown | Unknown |
| 162.92 | o2 | Unknown | Unknown |
| 164.909 | o2 | Halogenated VOCs | Chloral hydrate |
| 165.914 | o2 | Unknown | Unknown |
| 176.908 | o2 | Unknown | Unknown |
| 196.138 | o2 | Non-methane hydrocarbon | Unknown |
| 197.15 | h3o | Oxygenated VOCs | Unknown |
| 206.898 | o2 | Unknown | Unknown |
| 220.95 | o2 | Unknown | Unknown |
| 234.92 | h3o | Unknown | Unknown |
| 282.036 | o2 | Unknown | Unknown |
| 283.039 | o2 | Unknown | Unknown |

**Supplementary Figure 2.**
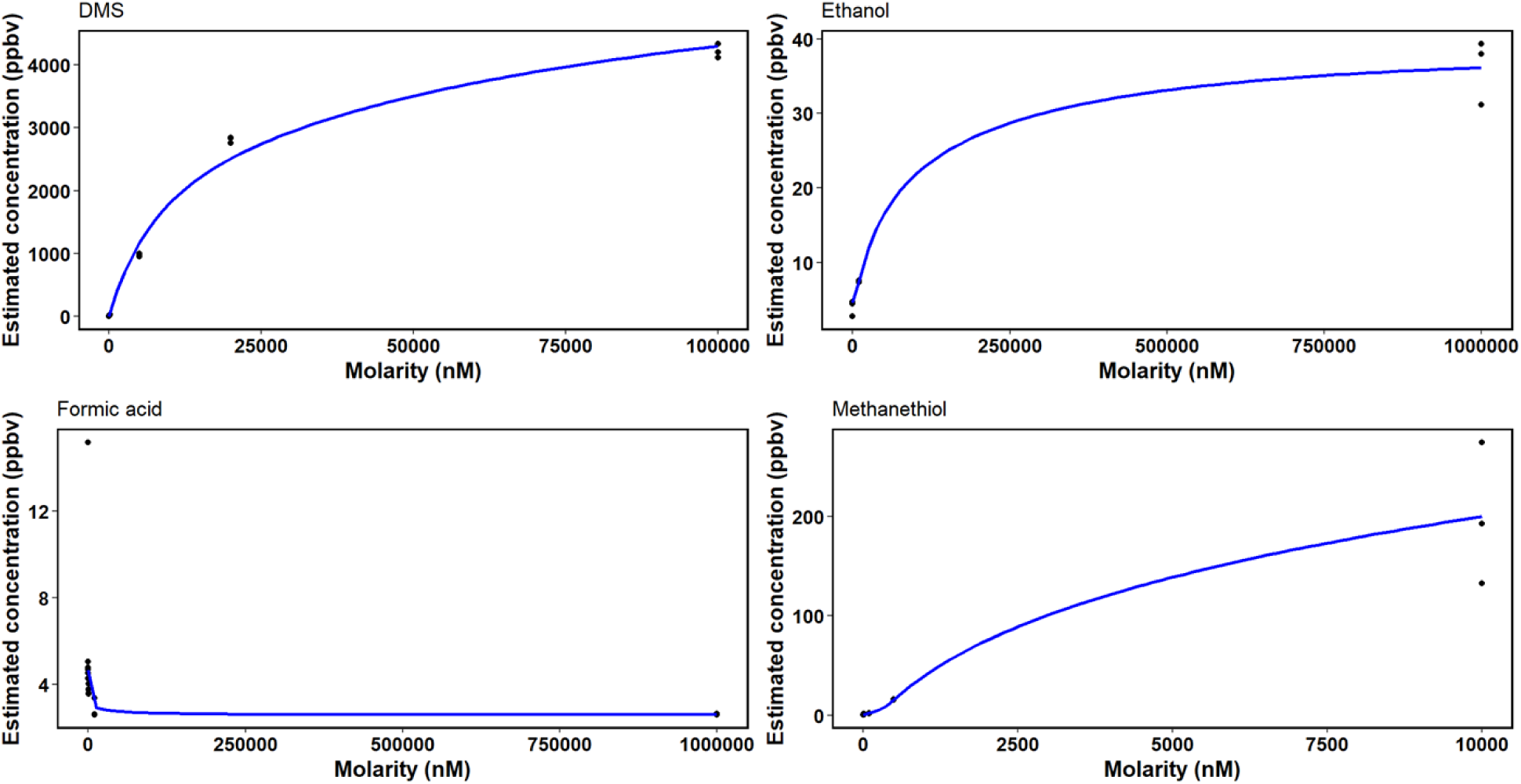
Standard curves of selected VOC compounds, comparing estimated concentration based on VOC signal strength with prepared known concentrations of DMS, Ethanol, Formic acid and Methanthiol to blank media.

**Supplementary Figure 3.**
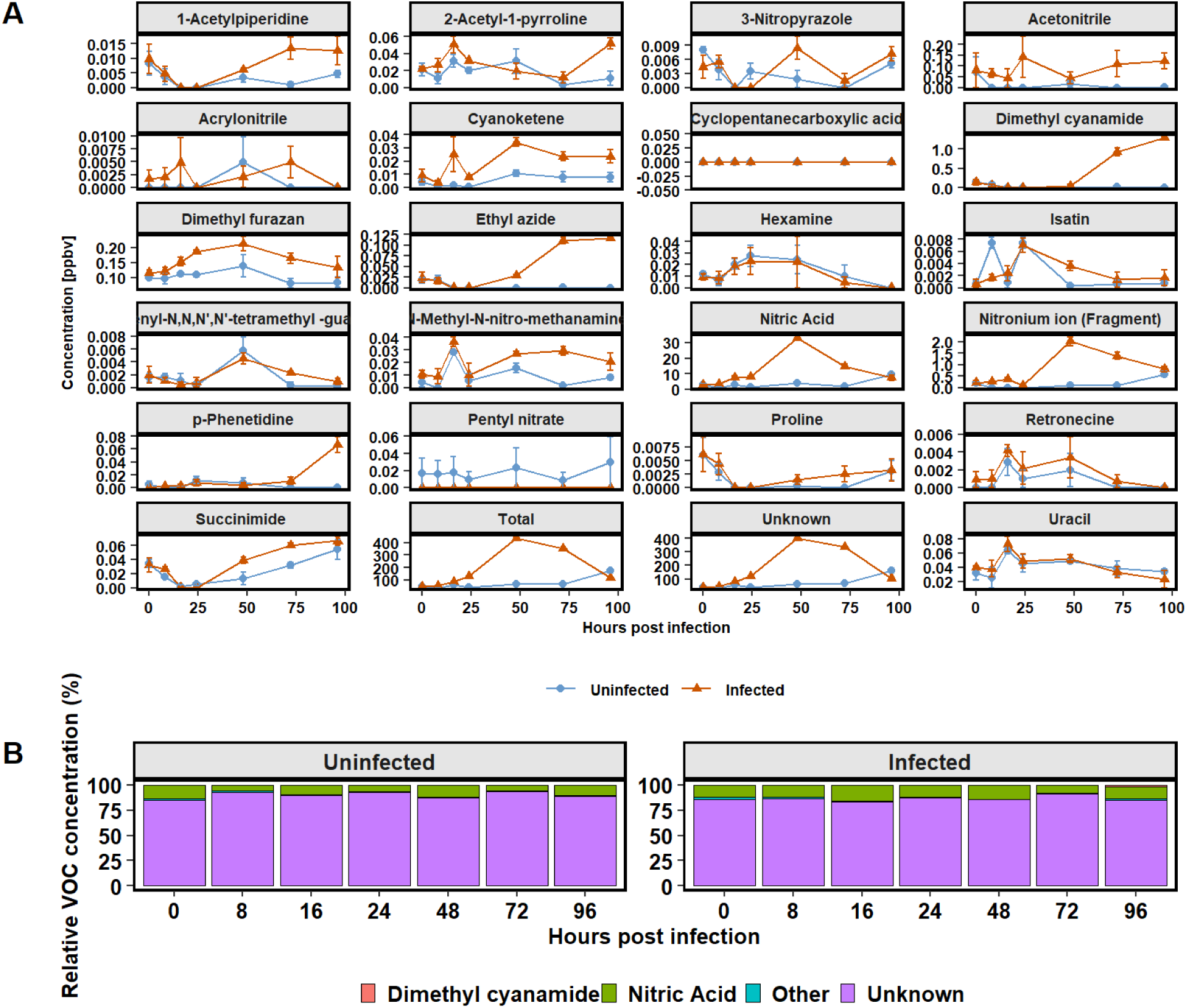
Estimated concentration of nitrogen-containing VOCs in infected and uninfected *G. huxleyi* cultures. (**A**) Mean concentration (ppbv) of all detected n-compound over the course of the experiments, with error bars indicating the standard error. (**B**) Relative contribution of compounds per timepoint, trace compounds (less than 1 % contribution to total) were grouped as “Other”. For the majority (83.3% to 93.0%) of N-VOCs the exact configuration of the compound could not be determined, due to structural ambiguity of identical molecular masses, thus were binned into “Unknown”.

**Supplementary Figure 4.**
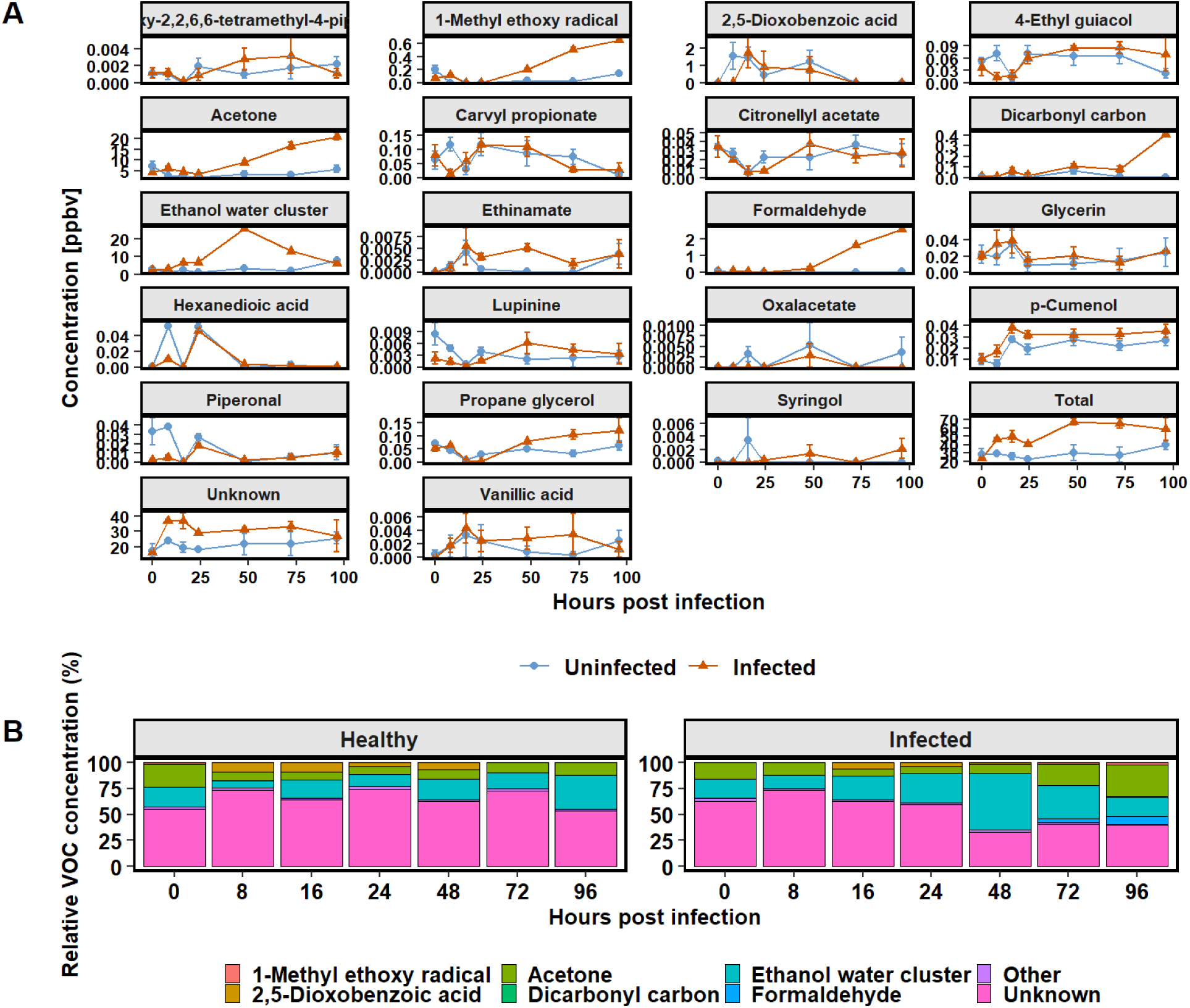
Estimated concentration of oxygenated VOCs in infected and uninfected *G. huxleyi* cultures. (**A**) Mean ppbv values of all detected compound of that VOC class over the course of the experiments, with error bars indicating the standard error. (**B**) Relative contribution of compounds per timepoint, trace compounds (less than 1 % contribution to total) were grouped as “Other”. For 36.5 % to 83.3 % of oxygenated VOCs the exact configuration of the compound could not be determined, due to structural ambiguity of identical molecular masses.

**Supplementary Figure 5.**
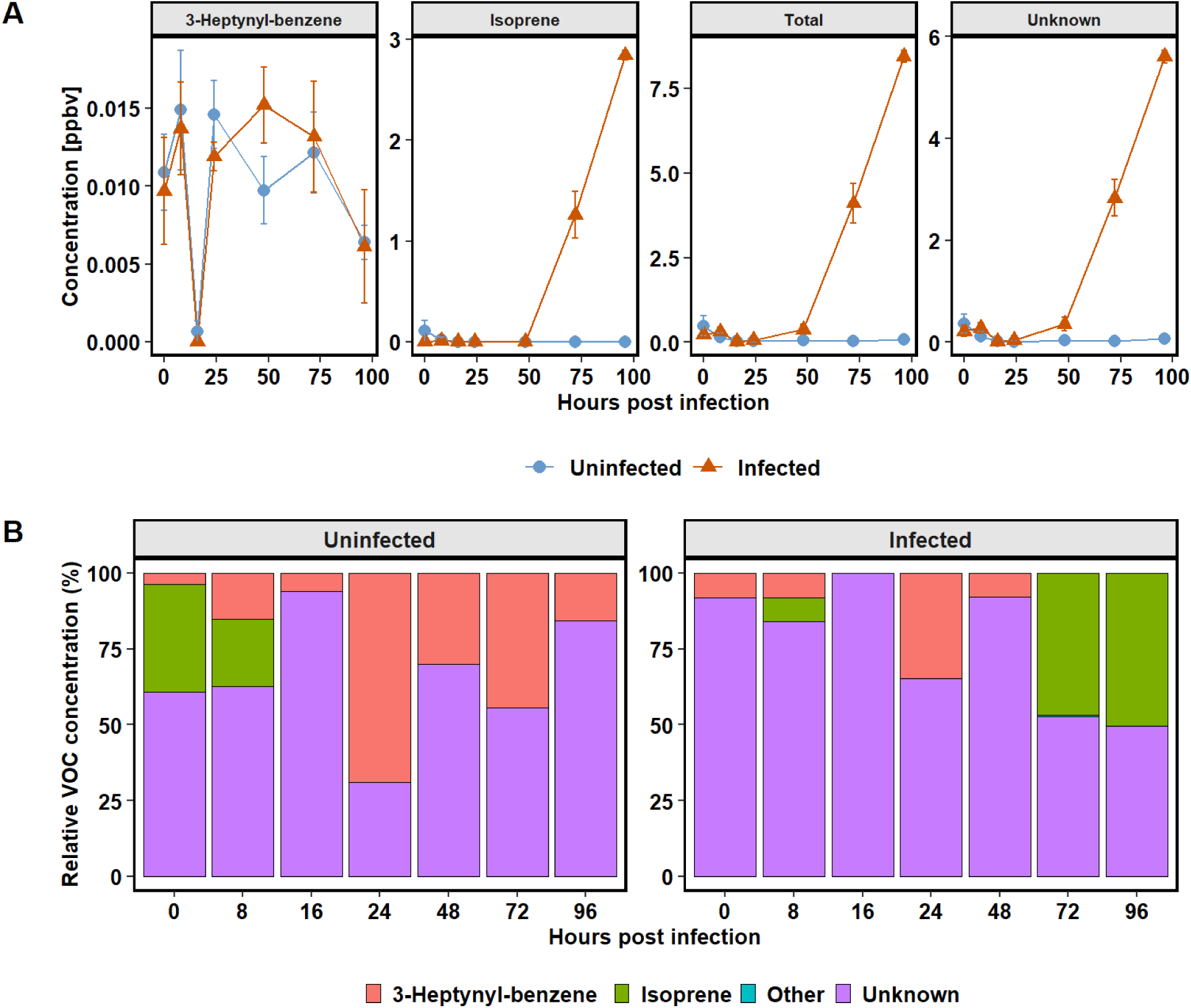
Estimated concentration of non-methane hydrocarbon VOCs in infected and uninfected *G. huxleyi* cultures. (**A**) Mean ppbv values of all detected compound categorized as hydrocarbons over the course of the experiments, with error bars indicating the standard error. (**B**) relative contribution of compounds per timepoint. Compounds binned into “Unknown” were due to structural ambiguity of identical molecular masses, which did not allow for identifying exact molecular configuration of those masses. Only a single hydrocarbon compound was identified (traces of 3-Heptynyl-benzene). Unlike other VOCs, hydrocarbons were only detected in the post-lysis phase in infected cultures, indicating non-biogenic release from cell debris.

**Supplementary Figure 6.**
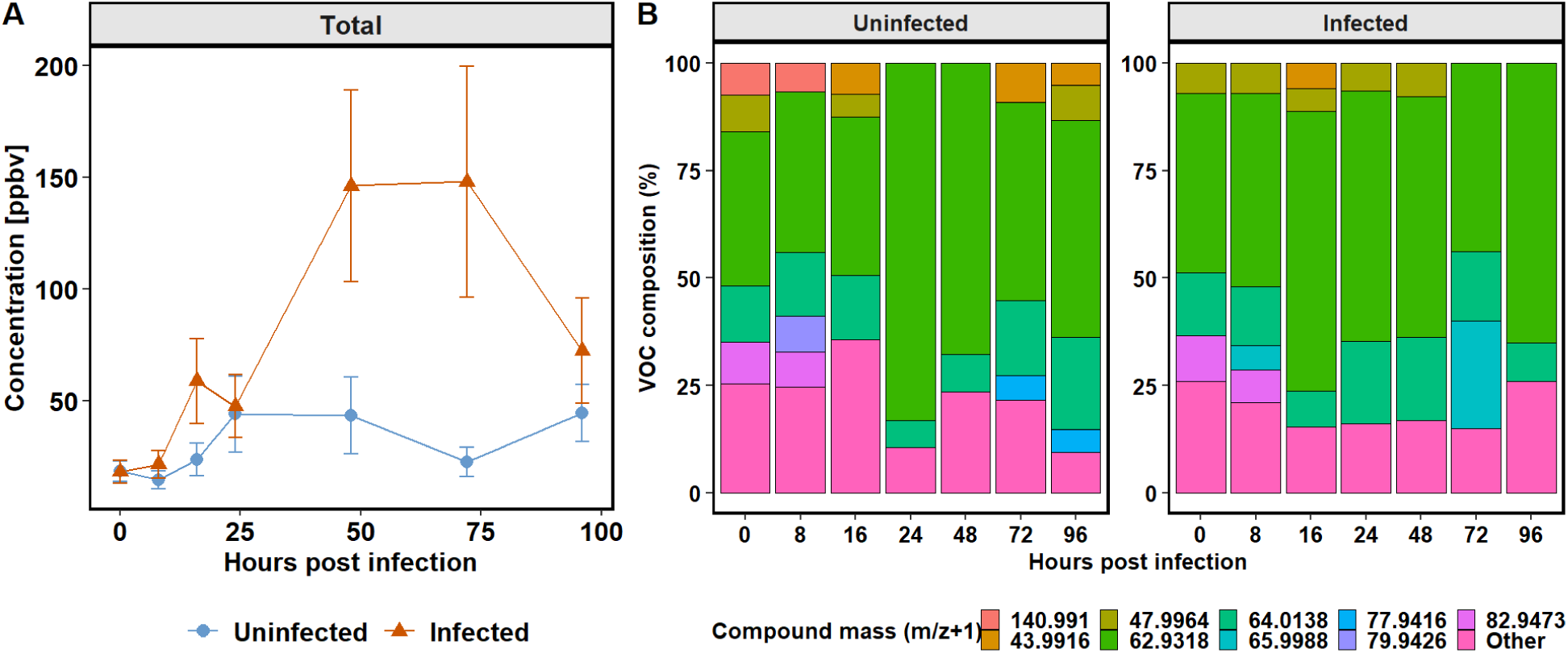
Estimated concentration of unidentified VOCs in infected and uninfected *G. huxleyi* cultures. (**A**) mean concentration (ppbv) that constitute >5% of unknown VOCs per timepoint; error bars indicating the standard error. (**B**) relative contribution (%) of those compounds per timepoint.

**Supplementary Figure 7.**
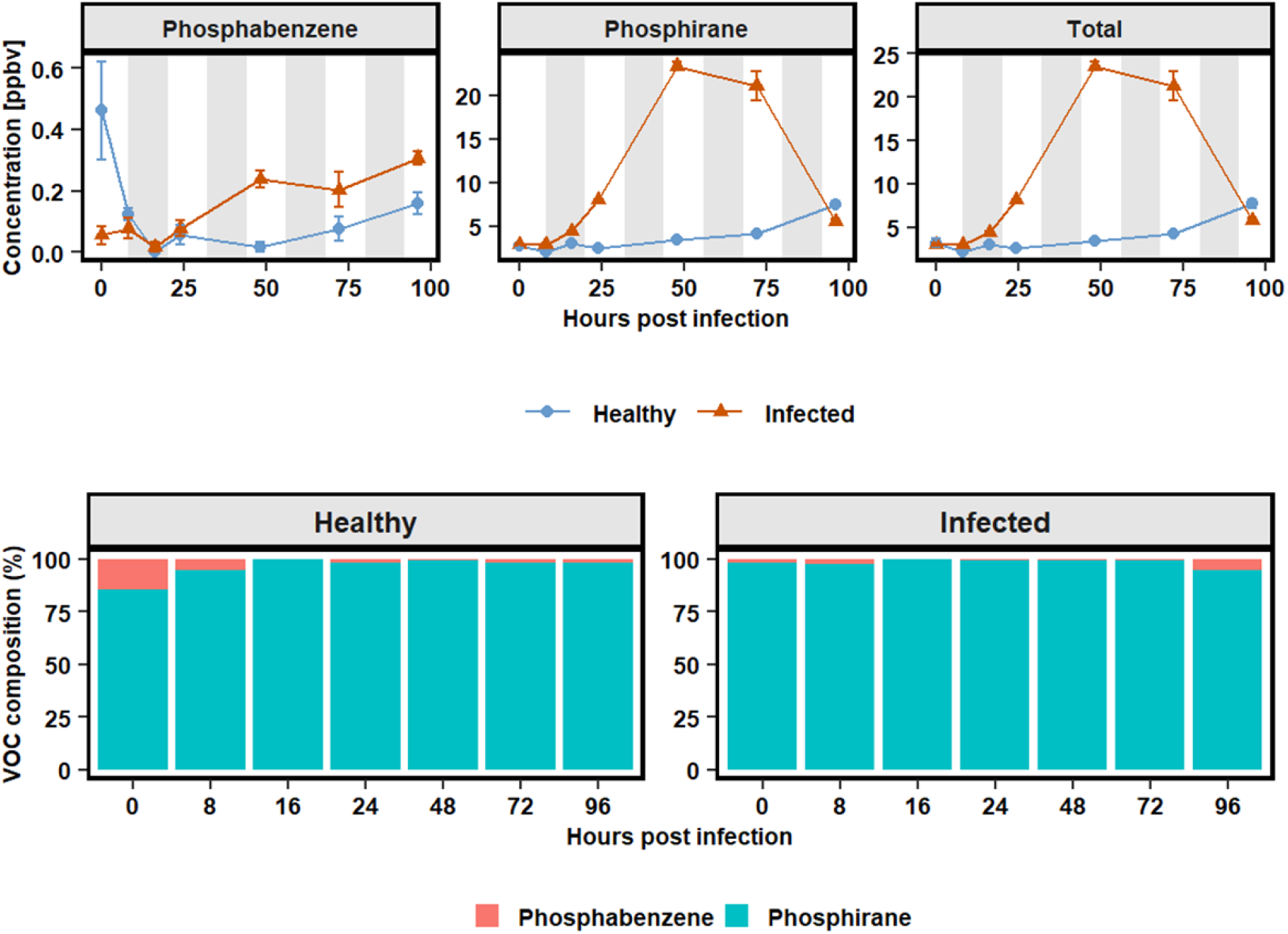
Estimated concentration of phosphor-containing VOCs in infected and uninfected *G. huxleyi* cultures. Top panels show the mean ppbv values of all detected compound categorized as p-containing, with error bars indicating the standard error. Stacked barplots display the relative contribution of compounds per timepoint.

## Notes

### Competing Interest Statement

The authors have declared no competing interest.

